# Genetic sex determination and sex-specific lifespan in tetrapods – evidence of a toxic Y effect

**DOI:** 10.1101/2020.03.09.983700

**Authors:** Zahida Sultanova, Philip A. Downing, Pau Carazo

## Abstract

Sex-specific lifespans are ubiquitous across the tree of life and exhibit broad taxonomic patterns that remain a puzzle, such as males living longer than females in birds and *vice versa* in mammals. The prevailing “unguarded-X” hypothesis (UXh) explains this by differential expression of recessive mutations in the X/Z chromosome of the heterogametic sex (e.g., females in birds and males in mammals), but has only received indirect support to date. An alternative hypothesis is that the accumulation of deleterious mutations and repetitive elements on the Y/W chromosome might lower the survival of the heterogametic sex (“toxic Y” hypothesis). Here, we report lower survival of the heterogametic relative to the homogametic sex across 138 species of birds, mammals, reptiles and amphibians, as expected if sex chromosomes shape sex-specific lifespans. We then analysed bird and mammal karyotypes and found that the relative sizes of the X and Z chromosomes are not associated with sex-specific lifespans, contrary to UXh predictions. In contrast, we found that Y size correlates negatively with male survival in mammals, where toxic Y effects are expected to be particularly strong. This suggests that small Y chromosomes benefit male lifespans. Our results confirm the role of sex chromosomes in explaining sex differences in lifespan, but indicate that, at least in mammals, this is better explained by “toxic Y” rather than UXh effects.

## MAIN

Sexual dimorphism in lifespan is widespread across the tree of life, including among tetrapods [1]. Which sex lives longer varies considerably in both direction and magnitude. For example, females live three times longer than males in the brown antechinus, a small marsupial, while males are twice as likely as females to survive from one year to the next in Arabian babblers, a passerine bird [2, 3]. In humans, the lifespan gap is estimated to be 4.8 years – female life expectancy is 74.7 years while male life expectancy is 69.9 years [4]. Understanding the dynamic nature of sex differences in lifespan across taxa remains an unsolved problem in evolutionary biology. To date, empirical studies have focused on adaptive hypotheses stemming from sex-specific differences in how natural selection operates on females and males [1, 5-8]. Despite their prominent role in explaining sex differences in ageing, adaptive processes seem unable to explain the observation that the homogametic sex tends to live longer than the heterogametic sex across a wide taxonomic range [1, 9].

There are at least two reasons why sex chromosomes should directly contribute to the evolution of different female and male lifespans [9, 10]. First, the unguarded X hypothesis (UXh) predicts increased mortality of the heterogametic sex due to the expression of deleterious recessive mutations that accumulate in the non-recombining parts of the X (or Z) chromosome [10]. Because these mutations are masked in the homogametic sex, males are predicted to live longer than females in ZW systems (where males are the homogametic sex), and females longer than males in XY systems (where females are the homogametic sex). This idea has received indirect support in the finding that adult sex-ratios are typically female-biased in tetrapods with XY systems, but male-biased in taxa with ZW systems, as expected if biased adult sex ratios result from sex-specific mortality [8]. A recent study further shows that the heterogametic sex tends to exhibit higher mean/maximum lifespan across a wide taxonomic range, but phylogenetic signal and sexual selection could contribute to explain this relationship [10]. In addition, recent experimental evidence shows that “unguarding” the X chromosome in females erases the sex gap in lifespan in *Drosophila melanogaster* [11, 12] but see [13]. A second hypothesis focuses on the role of the heteromorphic Y (or W) chromosome. Following recombination suppression, the non-recombining regions of Y (or W) chromosomes tend to accumulate deleterious mutations through evolutionary time, via processes such as Muller’s ratchet and genetic hitchhiking [14, 15]. Recombination suppression also leads to an accumulation of repetitive DNA in the Y and W chromosomes [14, 15], and recent evidence has shown that repetitive DNA is disproportionally de-repressed (and hence mis-expressed) during ageing in *Drosophila melanogaster*. Moreover, this phenomenon influences chromatin integrity and gene expression profiles genome-wide [16], and is more acute in old males than in old females [17, 18]. Therefore, in *D. melanogaster* there is solid evidence of substantial “toxic Y” effects, via both the accumulation of deleterious mutations and repetitive DNA elements, resulting in increased mortality of the heterogametic sex [13, 14]. However, the role of the “toxic Y” hypothesis in explaining broad patterns of sex differences in ageing has yet to be addressed.

The UXh and the toxic Y hypothesis are not mutually exclusive as both predict that sex differences in lifespan result from increased mortality of the heterogametic sex. However, for the former this is due to deleterious mutations in the X (or Z) chromosome, while for the latter this is due to the toxic effects of the Y (or W). This difference makes it possible to examine specific predictions regarding the relationship between female/male lifespan and the relative size of the sex chromosomes (Figure 1). Namely, the UXh predicts that the sex gap in lifespan (i.e. female - male lifespan) will be positively associated with: *a*) the degree to which the X chromosome is larger than the Y chromosome (and vice versa with Z/W), because “unguarded” recessive mutations have to accumulate in the non-recombining regions of the X (or Z) chromosome, and the overall size of these regions increases with the size difference between the sex chromosomes (Figure 1A), and *b*) the relative size of the X chromosome with respect to the rest of the genome (or *vice versa* for the Z), because this provides a measure of how much variability in lifespan we expect the X (or Z) chromosome to explain (Figure 1B). For example, X-linked effects are expected to be significant in *Drosophila melanogaster*, where the X chromosome constitutes ∼20% of the genome [19], but relatively minor in polar bears (*Ursus maritimus*), where the X chromosome is < 5% of the genome [20]. In contrast, toxic Y (or W) effects depend exclusively on the size of the non-recombining region in the Y (or W) chromosomes. Thus, toxic Y effects specifically predict lower male survival with increasing size of the Y (or W) chromosome relative to the autosomes (Figure 1C).

**Figure 1.**
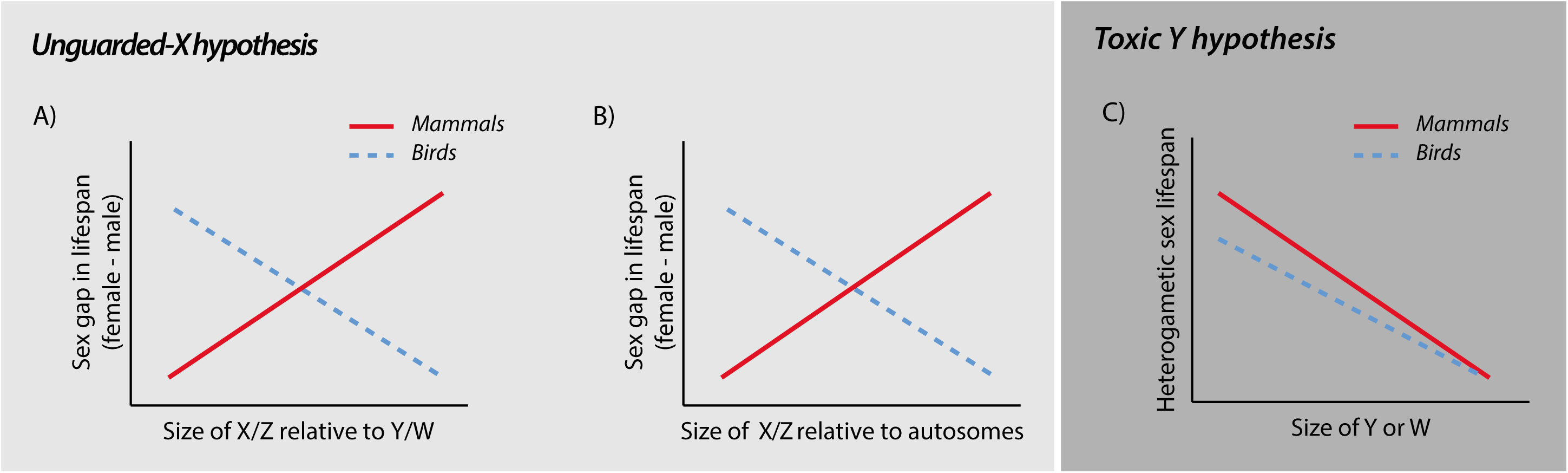
The unguarded X hypothesis predicts a positive relationship between the lifespan gap (i.e. female – male lifespan) and the size of the X relative to both the Y chromosome (A) and to the autosomes (B), and vice versa in the case of the Z chromosome. The toxic Y hypothesis, on the other hand, predicts a direct negative relationship between the size of Y and W chromosomes and the lifespan of the heterogametic sex (C). Toxic Y/W effects are expected to be stronger in mammals than in birds (e.g. because of the lower effective population size of Y compared with W).

We first collected data on sex-specific survival across 138 species of birds, mammals, reptiles and amphibians, representing 6 independent origins of XY systems and 6 independent origins of ZW system, and used phylogenetic meta-analytic models to test the general prediction that females are longer lived than males in XY systems and *vice versa* in ZW systems. To tease apart whether differences in survival between the sexes are driven by unguarded X or toxic Y effects, we then collected published karyotype data for 31 mammal and 15 bird species – the number for which we also had data on sex differences in survival. We focused on birds and mammals as we needed substantial variation in the sizes of sex chromosomes across species and this was lacking for amphibians and reptiles. We used this data to examine the following predictions. If the UXh contributes to the evolution of sex-specific lifespans, we expect: *i*) female mammals to have increasingly higher survival than males as the size of X relative to Y increases and as the size of X relative to the autosomes increases and *ii*) male birds to have increasingly higher survival than females as the size of Z relative to W increases and as the size of Z relative to the autosomes. If toxic Y effects contribute to the evolution of sex-specific lifespans, we expect: *i*) female mammals to have increasingly higher survival than males as the size of Y relative to the autosomes increases and *ii*) male birds to have increasingly higher survival than females as the size of W relative to the autosomes increases.

## RESULTS

### Sex differences in lifespan and genetic sex determination system across tetrapods

Females lived longer than males on average in XY systems while males lived longer than females on average in ZW systems (parameter estimate [*ß*] = −0.13, 95% Credible Interval [CI] = −0.20 to −0.06; *N*_*effect sizes*_ = 255; *N*_*species*_ = 138; Figure 2). The effect of the genetic sex determination system on sex differences in lifespan was independent of phylogeny, which explained 33% of the variance in sex-specific lifespans across species, and sexual size dimorphism (*ß* = 0.09, CI = −0.05 to 0.18; *N*_*effect sizes*_ = 255; *N*_*species*_ = 138), suggesting that differences in lifespan between the sexes are not due to overall differences in sexual selection. Considering each origin separately, in 6/6 XY systems females lived longer than males (in only one of these was the difference significant) and in 4/6 ZW systems males lived longer than females (in none of these was the difference significant)

**Figure 2.**
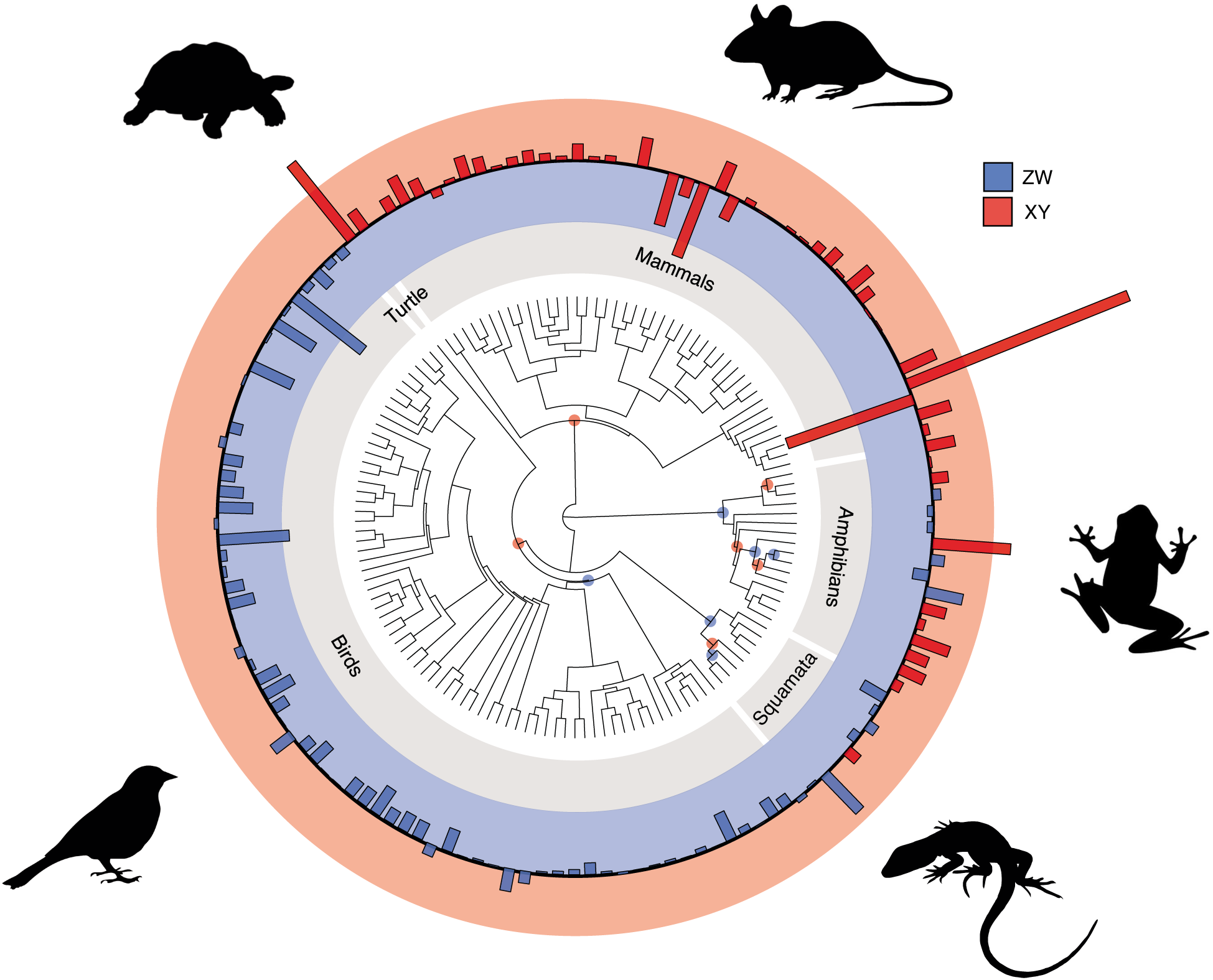
Sex differences in lifespan across 138 species of birds, mammals, reptiles and amphibians. A statistical effect size of the ratio of female and male lifespan (*lnR*) is plotted for each species. Bars above the black line (the red ring) indicate that females live longer than males, while bars below the black line (the blue ring) indicate that males live longer than females. ZW systems are coloured in blue and XY systems are coloured in red. The blue points in the phylogeny show the 6 independent origins of ZW systems and the red circles show the 6 independent of XY systems in our sample of species.

### Sex differences in lifespan and the difference in size between the sex chromosomes

The relative size of the X vs. Y chromosomes across mammals was not associated with sex-differences in lifespan (*ß* = −0.01, CI = −0.06 to 0.03; *N*_*effect sizes*_ = 55; *N*_*species*_ = 21; Figure 3a). In fact, the largest sex-differences in survival in mammals occur in species where X and Y appear to be similar in size (Figure 3a), as expected if large Y chromosomes have a negative effect on male survival rather than large X chromosomes. Sexual dimorphism in body size was also unrelated to lifespan differences between the sexes (*ß* = −0.02, CI = −0.09 to 0.04; *N*_*effect sizes*_ = 55; *N*_*species*_ = 21). Similarly, there was no relationship between sex-differences in lifespan and size differences between the Z and W chromosomes (*ß* = 0.02, CI = −0.07 to 0.10; *N*_*effect sizes*_ = 28; *N*_*species*_ = 14; Figure 3b) or sexual dimorphism in body size (*ß* = 0.02, CI = −0.04 to 0.10; *N*_*effect sizes*_ = 28; *N*_*species*_ = 14; Figure 3b) across birds.

**Figure 3.**
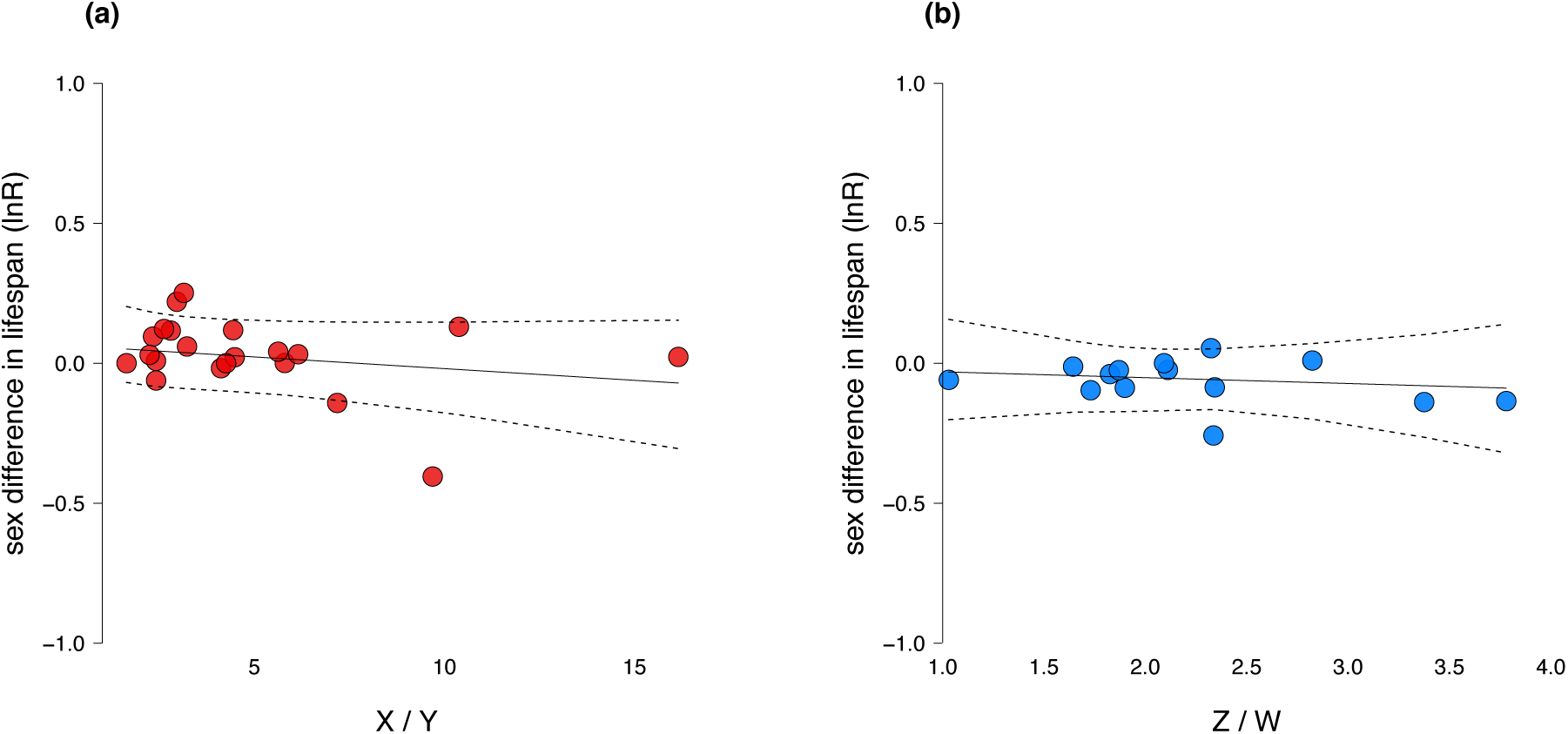
Sex differences in lifespan and the difference in size between the sex chromosomes in mammals (**A**) and birds (**B**). Species averages of effect sizes are plotted, with slopes and credible intervals estimated from phylogenetic meta-analytic models.

### Sex differences in lifespan and the relative sizes of the sex chromosomes

Across mammals, both the size of the X chromosome relative to the autosomes and the size of the Y chromosome relative to the autosomes were positively associated with sex differences in lifespan (X/autosomes: *ß* = 0.05, CI = 0.01 to 0.09; *N*_*effect sizes*_ = 65; *N*_*species*_ = 32; Figure 4a; Y/autosomes: *ß* = 0.05, CI = 0.00 to 0.11; *N*_*effect sizes*_ = 50; *N*_*species*_ = 20; Figure 4b). The larger the X and Y chromosomes relative to the autosomes, the larger the sex difference in lifespan, with females being increasingly long-lived compared with males. This, however, would be expected if the relative sizes of the X and Y chromosomes are themselves correlated. When modelling the effects of the relative sizes of the X and Y chromosomes together, the relationship between relative X chromosome size and sex differences in lifespan disappeared (*ß* = 0.01, CI = −0.05 to 0.06; *N*_*effect sizes*_ = 50; *N*_*species*_ = 20) while the relationship between relative Y chromosome size and sex differences in lifespan remained, although it was not statistically significant (*ß* = 0.04, CI = −0.01 to 0.11; *N*_*effect sizes*_ = 50; *N*_*species*_ = 20). Across birds, neither the size of the Z or W chromosomes relative to the autosomes were associated with sex differences in lifespan (Z/autosomes: *ß* = 0.01, CI = −0.07 to 0.07; *N*_*effect sizes*_ = 29; *N*_*species*_ = 15; Figure 4c; W/autosomes: *ß* = 0.00, CI = −0.09 to 0.07; *N*_*effect sizes*_ = 28; *N*_*species*_ = 14; Figure 4d).

**Figure 4.**
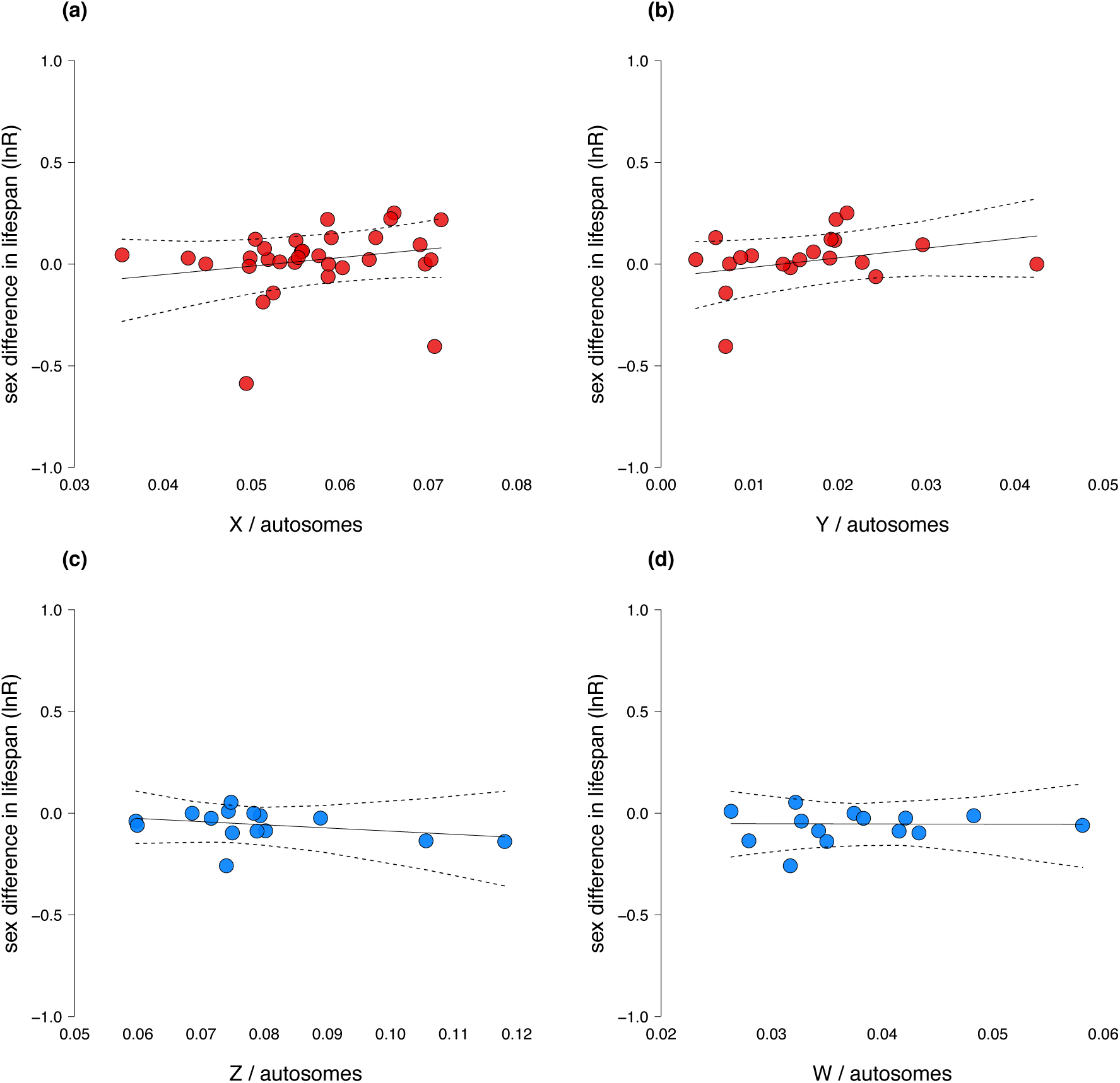
Sex differences in lifespan and the relative sizes of the sex chromosomes in mammals and birds. Sex differences in lifespan and the size of X relative to the autosomes (**A**) and Y relative to the autosomes in mammals (**B**). Sex differences in lifespan and the size of Z relative to the autosomes (**C**) and W relative to the autosomes in birds (**B**). Species averages of effect sizes are plotted with slopes and credible intervals estimated from phylogenetic meta-analytic models.

### Sex-specific lifespan and the relative sizes of the sex chromosomes

Male mammals with high rates of annual survival tended to have small Y chromosomes relative to the autosomes whereas males with low annual survival have relatively large Y chromosomes (*ß* = −0.72, CI = −1.19 to −0.26; *N*_*observations*_ = 50; *N*_*species*_ = 20; Figure 5a). This was independent of the strength of sexual size dimorphism (*ß* = −0.23, CI = −0.87 to 0.25; *N*_*observations*_ = 50; *N*_*species*_ = 20), a proxy for the strength of sexual selection, and differences in body size between species (*ß* = 0.35, CI = −0.01 to 0.76; *N*_*observations*_ = 50; *N*_*species*_ = 20). Moreover, male annual survival rates in mammals were not associated with the size of X chromosomes relative to autosomes (*ß* = −0.10, CI = −0.50 to 0.31; *N*_*observations*_ = 65; *N*_*species*_ = 32; Figure S1a) or with size differences between the X and Y chromosomes (*ß* = 0.17, CI = - to 0.72; *N*_*observations*_ = 55; *N*_*species*_ = 21; Figure S1b). In contrast, rates of female annual survival across bird species were not associated with the sizes of W chromosomes relative to autosomes (W/autosomes: *ß* = 0.13, CI = −0.51 to 0.67; *N*_*observations*_ = 28; *N*_*species*_ = 14; Figure 5b). Similarly, female annual survival in birds was not associated with the size of Z chromosomes relative to the autosomes (Z/autosomes: *ß* = −0.05, CI = −0.39 to 0.37; *N*_*observations*_ = 29; *N*_*species*_ = 15; Figure S1c) or with differences in size between the Z and W chromosomes (*ß* = −0.15, CI = −0.63 to 0.44; *N*_*observations*_ = 28; *N*_*species*_ = 14; Figure S1d).

**Figure 5.**
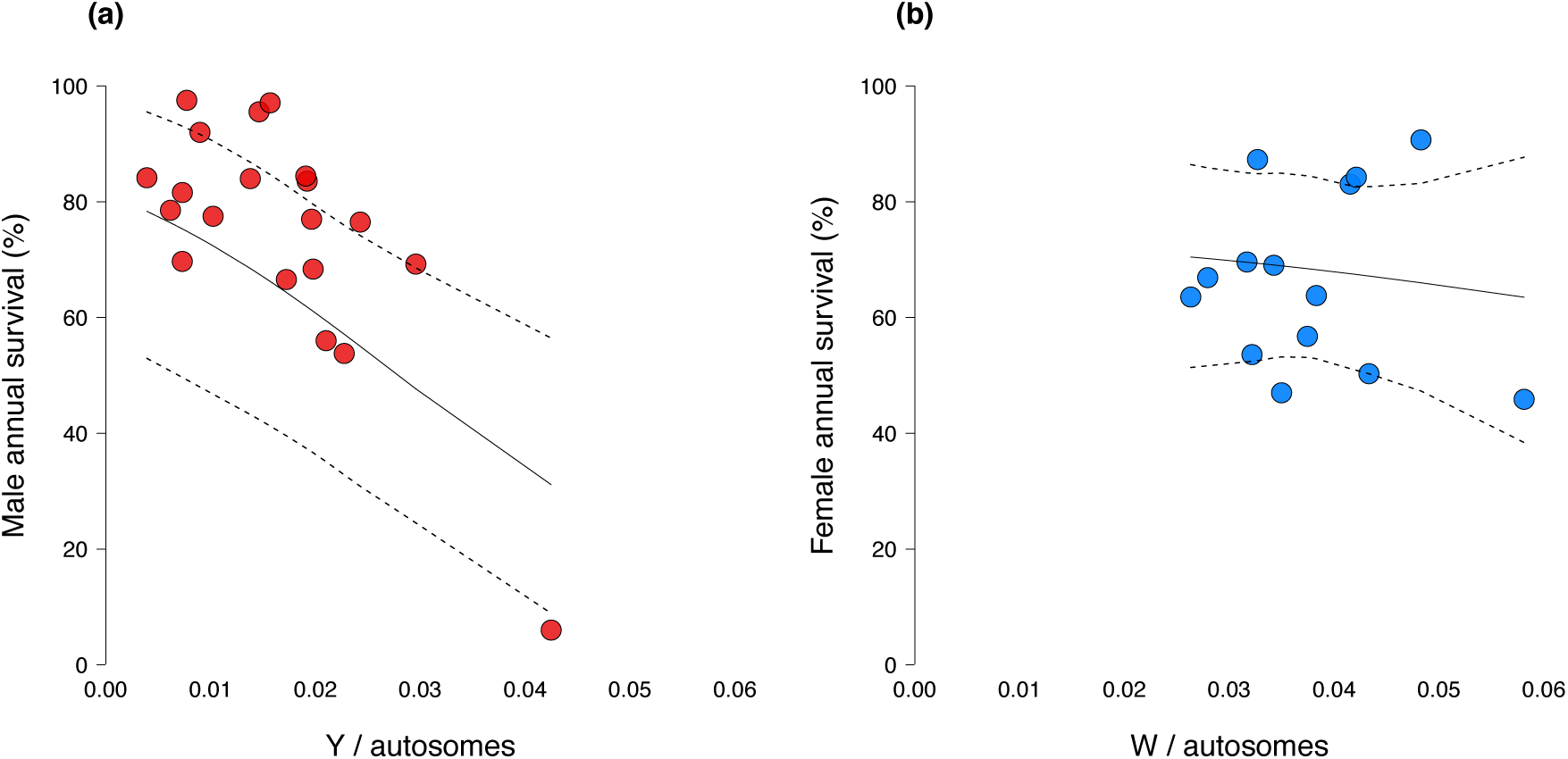
Sex-specific annual survival in relation to the relative size of the heteromorphic sex chromosome. Male annual survival and the relative size of the Y chromosome in mammals (**A**). Female annual survival and the relative size of the W chromosome in birds (**B**). Species averages of annual survival estimates are plotted with slopes and credible intervals estimated from phylogenetic mixed models.

## DISCUSSION

In this paper, we show that there is a clear link between genetic sex determination system and the sex gap in lifespan. Across 138 species of vertebrates reflecting 6 independent origins of XY and ZW systems, females survived longer than males on average in XY systems, while males survived longer than females on average in ZW systems. Previously, Pipoly et al. [9] identified female-biased adult sex ratios in taxa with XY systems and male-biased adult sex ratios in taxa with ZW systems. Along the same line, Xirocostas et al. [10] recently found that mean/maximum lifespan is shorter in the heterogametic sex across the tree of life, although the latter could be at least partly explained by the effects of phylogenetic signal and/or sexual selection. Our results build on these studies by showing that, in vertebrates, the relationship between sex differences in survival and sex determination system remains after accounting for both phylogenetic signal (which in our sample explained ∼33% of the variance in the sex gap in lifespan) and sexual size dimorphism (a proxy for sexual selection). Furthermore, our results suggest that this relationship is better explained by “toxic Y” rather than UXh effects, the prevailing hypothesis to date.

The most commonly cited hypothesis to explain broad patterns of sex-specific lifespan across taxa is the UXh hypothesis put forward by Trivers [21]. Recent work reporting a link between sex determination systems and different proxies for sex-specific lifespan have been interpreted as supporting this hypothesis [9, 10]. Here, we specifically tested for the UXh by asking whether there is a correlation between the sex-gap in survival and the relative size of the sex chromosomes (X/Z relative to Y/W and autosomes; Figure 1). We found limited evidence in its support (Figures 3 & 4). Despite its long history and intuitive appeal, our results suggest that the UXh may in fact not be a fundamental driver of sex-specific mortality. Since it was first formulated, more than three decades ago, support for the UXh has been scant and indirect [1, 9, 10, 12] and, while this may be due to the inherent difficulties in testing this hypothesis, there are also frequently overlooked reasons to doubt it plays a major role in explaining sex-specific lifespans. For example, the fact that there is likely to be strong selection against big effect X-linked recessive mutations in nature [22], meaning that any mutations that do accumulate on the recombining chromosome should have relatively minor effects. Of course, it is possible the lack of evidence in favour of the UXh hypothesis is simply due to a lack of statistical power in our study, given the relatively scarce karyotypic data generally available for vertebrates. We thus suggest that future analysis with more exhaustive datasets should aim to replicate our analysis. We note, however, that this same lack of power should affect our ability to detect “toxic Y” effects, a recent and until now empirically untested alternative that fits with the findings reported here.

According to theory, toxic Y effects result in reduced survival of the heterogametic sex due to the accumulation of both deleterious mutations and repetitive DNA elements in the Y and W sex chromosomes [14-18]. Two main predictions arise. First, that genetic sex determination systems predict the sex-gap in lifespan so that the heterogametic sex tends to live longer than the homogametic sex across a wide range of taxa, as reported here and in a recent study [10]. Second, that the size of the Y and W sex chromosomes inversely predict the lifespan of the heterogametic sex in XY and ZW systems respectively. Our results show that there is indeed a negative correlation between the relative size of the Y chromosome and male lifespan in mammals, although we did not find this effect in birds. However, a stronger toxic Y effect is expected in XY than in ZW systems because the effective population size of the Y chromosome is smaller than the W chromosome. This is due both to higher variance in male (than female) reproductive success and to Y chromosomes accumulating more mutations than W chromosomes because the male germ line undergoes more cell divisions than the female germ line [23]. This, in turn, makes degeneration of the Y chromosome (and hence accumulation of repetitive DNA) more likely [15]. For example, the mammalian Y chromosome is known to be significantly enriched in repetitive DNA compared to the W chromosome in birds [14, 15, 24, 25]. Available evidence also suggests that non-recombining regions are larger in mammalian than in bird sex chromosomes, which also seem to exhibit less variability in the Z/W (vs. mammalian X/Y) size ratio [14, 15, 24, 25]. In agreement with this, the X to Y chromosome ratio was twice as high on average than the Z to W chromosome ratio in our dataset (X to Y estimate = 4.96 ± 0.60 (se), *N*_*species*_ = 21; Z to W estimate = 2.23 ± 0.74, *N*_*species*_ = 14). This is not a sampling artefact as evidence from the karyotypes of 200 bird species shows that the Z to W chromosome ratio does not extend beyond the limit we detected [25].

An exciting possibility is that cyto-nuclear interactions may contribute to explain marked “toxic Y” effects in mammals, but not in birds. Given that Y chromosomes are not inherited along with mitochondrial DNA (and other cytoplasmic products), there is less scope for cyto-nuclear co-evolution in males vs. females with X/Y genetic determination systems (but see [26]). This is not the case in males of species with ZW sex-determination systems because in these species females are the heterogametic sex, and hence copies of both sex chromosomes are inherited maternally along with the cytoplasm. Indeed, recent evidence shows that cyto-nuclear interactions have sex-specific lifespan effects in *D. melanogaster* [27], which opens yet another exciting line of research for future studies. Although it is tempting to interpret our negative finding of a toxic W effect in line with the predictions above, this finding must be taken with caution for two methodological reasons: our sample size for birds was approximately half than that for mammals, and autosome size (on which many of our relative measures are based) is more difficult to estimate in birds than in mammals due to bird karyotypes often including a large number of micro-chromosomes that increased measurement error in our estimations of bird chromosome sizes.

Other complementary hypotheses that could indirectly explain a role of sex chromosomes in determining the sex gap in lifespan have to do with sexual selection and imperfect dosage compensation. First, sexual selection frequently favours the accumulation of mutations that increase male reproductive success at the expense of male survival [7], but in both our study and in Pipoly et al. [9] correlations between sex determination systems and survival/adult sex ratios were independent of sexual size dimorphism, a commonly used proxy for sexual selection intensity. Second, imperfect dosage compensation is deleterious for the heterogametic sex, explaining why this sex is short-lived relative to the homogametic sex [28]. However, imperfect dosage compensation predicts that the sex gap in lifespan should be related to the size difference between the two sex chromosomes (X relative to Y and Z relative to W) because the size difference will be proportional to the imbalance in gene dose between males and females, and hence the degree of dosage compensation [28]. Our results show, however, that this was not the case (Fig 3).

To conclude, we report compelling evidence of a link between sex determination systems and the sex-gap in survival across vertebrates, which along with recent evidence [10] strongly suggests that sex chromosomes play a role in understanding broad patterns of sexual differences in lifespan. Our data suggest that “toxic Y” effects, rather than unguarded X effects explain this link. Future research should aim to explicitly test for toxic Y effects by means of direct empirical manipulations [16, 18] and by expanding our comparative framework to other taxa (e.g. invertebrates).

## METHODS

### Data Collection

To detect unguarded X and toxic Y effects using comparative analyses, we required data on sex-specific lifespans across multiple independent origins of XY (male heterogametic) and ZW (female heterogametic) systems as well as variation across species in the extent of recombination suppression between the sex chromosomes. We therefore limited our search to tetrapods (amphibians, reptiles, birds and mammals) with XY or ZW systems because of the abundance of species that have been relatively well studied with respect to sex-specific lifespans (or proxies thereof, see below). We excluded fish because we did not find enough concurrent data on both sex-specific lifespan and genetic sex determination system.

#### Sex differences in lifespan

We used sex-specific survival rates and mean ages as proxies for sex differences in lifespan to maximise available data across multiple independent origins of XY and ZW systems. Average lifespans are difficult to measure in wild vertebrate populations and life-tables (from which adult life expectancy can be calculated) have been reported for relatively few species, primarily mammals and birds [2, 29]. Both survival and mean age are directly related to lifespan: life history theory predicts a longer lifespan when extrinsic mortality is low and thus survival is high while mean age will be lower in populations with a higher proportion of young individuals. Furthermore, survival and mean age are typically estimated using mark-recapture methods and so have the advantage of being reported with measurement error, which can be incorporated into statistical analyses. We excluded maximum lifespan as a proxy for average lifespan because it is strongly dependent on sampling effort, it is reported without error (by definition) and currently available data come from a mixture of wild and captive populations frequently based on very few individuals [30].

We used Scopus and Web of Science (WoS) to identify studies reporting sex-specific survival rates and mean ages. First, we used the following topic search term: “*(male or female) AND mark-recapture*” (studies published up to 31/12/2018). This returned 1647 studies from WoS and 2227 studies for Scopus. We also conducted backward and forward citation searches on the following review papers of survival rates [31-36]. This returned 168, 42, 118, 91, 8 and 31 studies from each review respectively. This search strategy primarily returned studies with suitable data on birds and mammals. To increase the number of amphibians and reptiles in our sample, we therefore conducted two additional searches in WoS and Scopus using the following topic search terms “*mean age OR average ag AND male AND female AND amphibian OR reptile*” and “*skeletochronology AND male AND female*”. The first search terms returned 25 studies from WoS and 32 studies from Scopus and the second search terms returned 205 studies from WoS and 207 studies from Scopus. We included skeletochronology in our second search term because this is a widely-used technique for determining mean age in reptiles and amphibians and is typically reported with error for females and males. We screened the title of each study in our sample (*N* = 4801 studies) and the abstracts of those which appeared to contain relevant data and identified 157 studies with suitable data (female and male survival / mean age reported with error) and that matched with the reptile and amphibians with known genetic sex-determination systems (see below).

#### Genetic sex-determination system

We collected data on the genetic sex determination system for each species in our database using the tree of sex database [37]. We complemented this with individual searches for species for which we had data on sex-specific lifespan (i.e. survival/mean age) but that did not appear in the tree of sex database; using the keywords “species name” AND “karyotype” OR “sex chromosome” in Google Scholar, Scopus and WoS and then examining the abstracts in search for data on the genetic determination system (see SM). All birds were assigned as ZW (*N*_*species*_ = 69) and all mammals as XY (*N*_*species*_ = 45). We assigned six amphibian and seven reptile species as ZW and ten amphibian and two reptile species as XY. We estimated the number of independent origins of XY and ZW systems in our sample of species using data from published sources (Evans et al. 2012 Chapter 18 in Polyploidy and Genome Evolution). There were 2 independent XY origins and 2 independent ZW origins in our sample of reptiles and 3 independent XY origins and 3 independent ZW origins in amphibians. Therefore, in total, our sample included six XY origins and six ZW origins (see SM for details).

#### Karyotypes

We collected karyotype data for all available bird and mammal species for which we also had sex-specific survival/mean age data. We focused on birds and mammals as we needed variation in the sizes of the recombining and non-recombining chromosomes across species that share the same origin of a genetic sex determination system. This was not possible for amphibians and reptiles where each independent origin of XY or ZW systems included seven or fewer species. We began by searching 14 chromosome atlases [20, 38, 39] for the karyotypes of species for which we had data on sex differences in survival. We identified karyotype images for 22 mammal and 4 bird species. For the species that were not present in these atlases, we did an additional search using the keywords “*species name AND karyotype*” in Web of Science and Scopus. We then examining the content of each article for karyotype images. From these searches, we found karyotype images for a further 9 mammal and 11 bird species. In total, we found suitable data for 32 mammal and 15 bird species. For each of these species, we used the karyotype image to calculate three ratios: *i*) the ratio of X or Z to the rest of the chromosomes (X / autosomes or Z / autosomes), *ii*) the ratio of Y or W to the rest of the chromosomes (Y / autosomes or W / autosomes) and *iii*) the ratio of X to Y and W to Z (X / Y or Z / W). We used ImageJ [40] for calculating the relative size of the sex chromosomes from karyotype images (see SM).

#### Sexual size dimorphism

We used sexual size dimorphism as a proxy to control for the intensity of sexual selection by including this variable as a co-variate in our statistical models (see below). We aimed to collect data on female and male head to body length for mammals, amphibians and reptiles. However, sex-specific body lengths were not available for most of the birds and some of the mammals in our sample, in which cases we extracted data on female and male body mass instead (see Table S1). For mammals, we complied data from two online resources: http://www.arkive.org and http://eol.org. For the species that were not present in these databases, we did an additional search by using the keywords “*species name AND body size AND male AND female*” in WoS and Scopus, and then examining the whole content of the articles in search of male/female body length data. From this additional search, we found sexual size dimorphism data for X more mammal species. For amphibians and reptiles, the studies that we took the sex-specific mean age data from also provided information about sex-specific body size (see Table S1). For birds, sexual size dimorphism was taken from the Handbook of the Birds of the World [41]. Finally, we calculated sexual size dimorphism as the natural logarithm of the ratio of female to male body size: *ln (female value / male value)*.

#### Phylogenetic trees

We used the *rotl* R package, an interface to the Open Tree of Life [42], to estimate a phylogenetic tree of the relationships among species in our sample. Branch lengths were estimated using [43] method in the *APE* package in R, with each node height raised to the power of 0.5. This tree was used in the analysis of sex differences in lifespan across tetrapods (see below). For the analysis involving birds only, we downloaded a sample of 1300 phylogenies (out of 10 000) from the birdtree.org [44]. Similarly, for the analysis involving mammals only, we downloaded a sample of 1300 phylogenies (out of 10 000) from URL [45]. We calculated a phylogenetic covariance matrix (evolutionary distances between species) from each of these trees which were then used to account for dependencies due to shared evolutionary history in our statistical models.

### Effect size calculation

We compared female and male lifespans using an effect size which allows us to take a standardised measure of the magnitude of the statistical difference between females and males that is comparable between [46]. We used the natural logarithm of the response ratio:

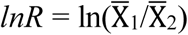

where 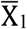 is either female mean age or female annual survival and 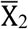 is either male mean age or male annual survival depending on which data were available for a given species. Positive values indicate that females live longer than males and negative values that males live longer than females. Each effect size was weighted by its sampling variance in our statistical models to account for differences in sampling effort between studies:

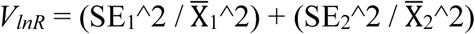

where SE_1_ is the standard error of the female value and SE_2_ is the standard error of the male value. In total, we obtained 255 effect sizes from 157 studies on 138 species across 6 independent origins of XY systems and 6 independent origins of ZW systems. There was significant between-study variance (τ^2^ = 0.01, *I*^2^ = 97.1%, Cochran’s *Q* = 8073.7, p < 0.001, *N*_*effect sizes*_ = 255). We detected no evidence of publication bias using Egger’s regression method (intercept = 0.00, p = 0.96). However, a trim and fill analysis suggested that 54 effect sizes were missing from our sample (Figure SX i.e. contour-enhanced funnel plot). There was no difference in mean *lnR* between studies reporting annual survival and those reporting mean age in XY or ZW systems (XY difference = −0.09, se = 0.06, p = 0.13, *N*_*effect sizes*_ = 120; ZW difference = −0.07, se = 0.05, p = 0.15, *N*_*effect sizes*_ = 135) suggesting that the proxy used to estimate sex differences in lifespan does not bias our effect sizes. We report heterogeneity statistics for each of the meta-analytic models described below in the supplementary materials, including the percentage of variation in *lnR* attributable to phylogenetic history and repeated observations made on the same species [47]. Full details are provided in the supplementary R code.

### Statistical models

We used the metafor [48] and MCMCglmm [49] R packages for model fitting when analysing *lnR*. Metafor uses restricted maximum likelihood for parameter estimation while MCMCglmm uses the Markov chain Monte Carlo (MCMC) method in a Bayesian framework. For analyses of non-Gaussian response variables (i.e. sex-specific annual survival rates, described below) we used MCMCglmm only which has greater flexibility when fitting models with non-normal distributions. We therefore report parameter estimates from MCMCglmm in the results section for consistency between analyses but show those from metafor (see SM). Parameters are reported as the posterior mode (*ß*) and 95 % credible interval (CI) of the posterior distribution of the Markov chain and significance is assessed by whether the CI includes zero (ref). For the MCMCglmm models, we used uniform priors for fixed effects and inverse-Wishart priors (v = 1 and nu = 0.002) for random effects and ran each model for 1,300,000 iterations with a burn in period of 300,000 and saving every 1000^th^ iteration of the chain. Full details including model diagnostics are provided in the supplementary R script.

#### I. Sex differences in lifespan and genetic sex determination system across tetrapods

To test whether females are longer-lived than males in XY systems and males are longer-lived than females in ZW systems we modelled *lnR* (treated as Gaussian) as a function of genetic sex determination system (2 level fixed effect: ZW or XY) with sexual size dimorphism included as a covariate (z transformed: mean = 0 and sd = 1) and the phylogenetic covariance matrix from the tetrapod phylogeny created using the *rotl* R package (see above) as a random effect. We also included a species-specific random effect to account for repeated measures made on the same species. Each effect size was weighted by its sampling variance, *V*_*lnR*_.

#### II. Sex differences in lifespan and the difference in size between the sex chromosomes

We modelled the relationship between sex differences in lifespan and the difference in size between the sex chromosomes (X vs. Y and Z vs. W) to test for an unguarded X effect. If recessive mutations accumulate in the non-recombining regions of X or Z chromosomes then the larger the size difference between the X and the Y (or between Z and W), the larger the non-recombining region, resulting in more recessive mutations and greater sex differences in lifespan. First, we modelled *lnR* (treated as Gaussian and weighted by sampling variance) as a function of X/Y (log and z transformed) in mammals. We included sexual size dimorphism as a covariate (z transformed) and a species identifier and a phylogenetic covariance matrix as random effects. We replaced the phylogenetic covariance matrix used in each model every 1000 iterations of the Markov chain with the next one in the sequence calculated from the 1300 phylogenies downloaded from URL. The values of the variance components and latent variables estimated using the previous phylogenetic covariance matrix were used as starting values for the next one in the sequence. This allowed us to incorporate uncertainty in the mammal phylogeny into our analyses. Note that this was not possible for the metafor analysis and parameters were calculated based on one randomly sampled phylogenetic covariance matrix. Next, we modelled *lnR* (treated as Gaussian and weighted by sampling variance) as a function of Z/W (log and z transformed) in birds. Sexual size dimorphism was included as a covariate (z transformed) and a phylogenetic covariance matrix and a species identifier were included as random effects. The phylogenetic covariance matrix used in each model was updated as described for the mammal analyses using the phylogenetic covariance matrices calculated from the 1300 phylogenies downloaded from birdtree.org.

#### III. Sex differences in lifespan and the relative sizes of the sex chromosomes

The UX hypothesis predicts that sex differences in lifespan should correlate with the size of the X (or Z) chromosome relative to the rest of the genome, as this ratio measures the potential impact of recessive mutations on survival. Similarly, the toxic Y hypothesis predicts that sex differences in lifespan should correlate with the size of the Y (or W) chromosome relative to the rest of the genome. First, we tested for UX effects by constructing two models. In mammals, we modelled *lnR* (treated as Gaussian and weighted by sampling variance) as a function of X/autosomes (z transformed) and sexual size dimorphism (z transformed) with a phylogenetic covariance (updated as described above) and a species identifier included as random effects. We repeated this model for birds, substituting X/autosomes for Z/autosomes. We then tested for toxic Y effects in mammals and birds by repeating the above models but replacing X/autosomes with Y/autosomes (z transformed) in the mammal analysis and Z/autosomes for W/autosomes (z transformed) in the bird analysis. All other fixed and random effects were the same. Finally, to tease apart toxic Y and unguarded X effects we modelled *lnR* as a function of both X/autosomes and Y/autosomes (both z transformed) in mammals and both Z/autosomes and W/autosomes (both z transformed) in birds. We included sexual size dimorphism as a covariate (z transformed) and a phylogenetic covariance matrix (updated as described above) and a species identifier as random effects in each model. Effect sizes were weighted by their sampling variance. The X/autosome and Y/autosome ratios were weakly correlated in mammals (r = 0.23, lwr = −0.23, upr = 0.61, p = 0.32)) and Z/autosome and W/autosome ratios were weakly correlated in birds (r = −0.31, lwr = −0.72, upr = 0.26, p = 0.29).

#### IV. Sex-specific survival and the sex chromosomes

The unguarded X and toxic Y hypotheses predict reduced survival of the heterogametic sex specifically. We tested this in mammals by comparing male annual survival rates (males are XY) across species in relation to *i*) X/Y (log and z transformed), *ii*) X/autosomes (z transformed) and *iii*) Y/autosomes (z transformed). Male annual survival was modelled using a binomial distribution (number alive vs. number dead) with a logit link function. In each of these three models we included male body mass (z and log transformed) and sexual size dimorphism (z transformed) as covariates to control for variation in male annual survival rates across species explained by differences in the strength of sexual selection and size differences. A phylogenetic covariance matrix (updated as described above) and a species level identifier were included as random effects in each model. Females are the heterogametic sex in birds, therefore we modelled female annual survival rates (using a binomial distribution with a logit link function) as a function of *i*) Z/W (log and z transformed), *ii*) Z/autosomes (z transformed) and *iii*) W/autosomes (z transformed). In each of these three models, we included female size as a covariate, to account for variation in female survival rates across species explained by differences in size, and a phylogenetic covariance matrix (updated as above) and a species level identifier were included as random effects.

## Acknowledgments

We thank Fernando Gonzalez Candelas and Mauro Santos for stimulating discussions about this study. PC was supported by research grants (CGL2014-58722-P and CGL2017-89052-P) from the Plan Nacional I+D+i by the Spanish Government, co-funded by the European Regional Development Fund, and by a Ramón y Cajal fellowship (RYC-2013-12998) also by the Spanish Government. ZS was supported by a PhD fellowship from the University of Valencia.

## Figures

**Supplementary Figure 1.**
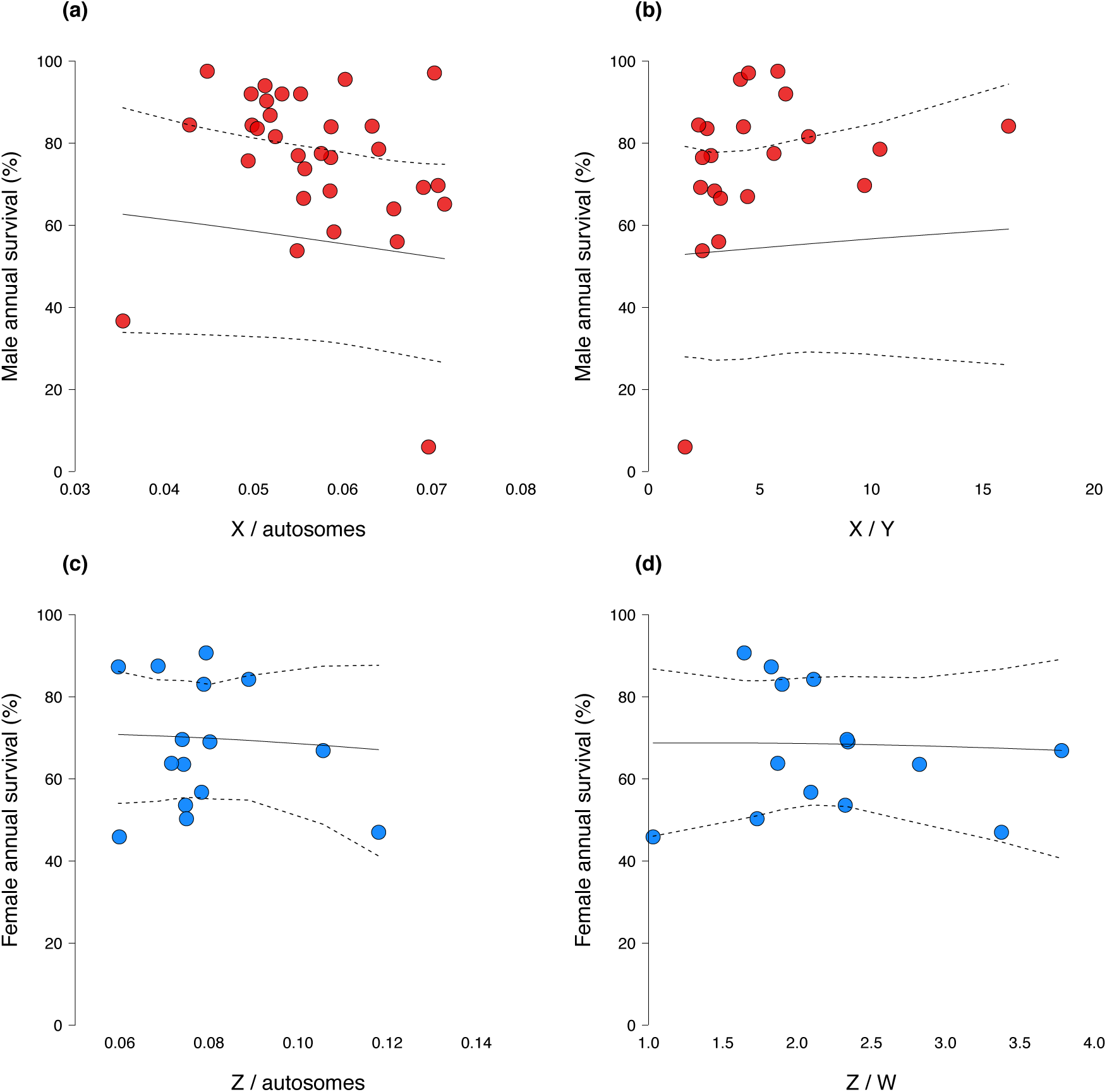
Sex-specific annual survival in relation to the relative size of the homomorphic sex chromosomes and to the difference in size between the sex chromosomes. Male annual survival and the relative size of the X chromosome (**A**) and the difference in size between X and Y (**B**) in mammals. Female annual survival and the relative size of the Z chromosome (**C**) and the difference in size between Z and W (**D**) in birds. Species averages of annual survival estimates are plotted with slopes and credible intervals estimated from phylogenetic mixed models.

